# Murine implantation chamber formation precedes natural and artificial decidualization

**DOI:** 10.64898/2025.12.30.697086

**Authors:** Harini Raghu Kumar, Noura Massri, Aishwarya V Bhurke, Akanksha Kapur, Pooja Gadhiya, Ripla Arora

## Abstract

During pregnancy uterine stromal cells undergo a mesenchymal to epithelial cell transition termed decidualization. In humans initiation of decidualization occurs in the absence of an embryo resulting in a need to identify embryo-independent molecular cues that can initiate decidualization. Although, similar to humans, decidualization in the mouse can be induced in the absence of an embryo, whether an implantation event is prerequisite for such decidualization is not known. In this study using different models of estrogen-dependent implantation, including natural (embryo) and artificial (sesame oil, agarose only beads, and Concanavalin A coated agarose beads) we determined that implantation chamber formation precedes decidualization. We show that focal stimuli, including the embryo, Concanavalin A coated bead, and oil droplets, induce V-shaped implantation chambers that lead to sub-epithelial PTGS2 expression and decidualization. Unfertilized eggs and uncoated agarose blue beads fail to form an implantation chamber and do not initiate decidualization. Further, we show that lectins that share sugar binding properties with Concanavalin A can also induce a V-shaped implantation chamber. Finally, using second harmonic generation we show that during decidualization collagen fibers spread radially away from the implantation chamber irrespective of the focal signal used for inducing the chamber. Thus, in the mouse artificial decidualization also initiates at the site of implantation chamber formation. These findings are critical when separating physical stimulus-dependent, embryo-dependent and embryo-independent mechanisms of decidualization that underlie a successful pregnancy.

## INTRODUCTION

Uterine architectural changes are key to embryo movement, embryo spacing, embryo-uterine interactions and implantation chamber formation in early mouse pregnancy (Flores et al., 2020; Madhavan et al., 2022). Attachment of the embryo to the uterine luminal epithelium is followed by dramatic changes in uterine mesenchymal (stromal) cell shape and secretory function defined as decidualization (Lee and DeMayo, 2004; Maurya et al., 2021). While initiation of decidualization in humans occurs every cycle during the progesterone rich secretory phase, decidualization in mice is strictly dependent on the presence of an embryo (Rodolfo et al., 2014). However, the relationship between embryo implantation and the onset of murine decidualization is unclear.

The search for a decidual stimulus is ongoing since the beginning of the 19^th^ century. Typical assays for inducing artificial decidualization or a deciduoma include increase in uterine weight and expression of alkaline phosphatase in the stroma (Finn and Hinchliffe, 1964). The earliest reports in 1937 suggested that electric current can induce decidualization (Krehbiel, 1937). It has also been well established that trauma to the uterine epithelium using a syringe needle during the receptive period can induce formation of a deciduoma (Lee et al., 2007). Glass beads, paraffin beads and tumor cells injected into the uterus during pseudopregnancy also induce a decidual reaction (Blandau, 1949; Rous, 1911; Wilson, 1963). These studies suggested that focal objects can induce decidua, leading to the hypothesis that focal pressure is required for formation of decidua. However, eggs present in pseudopregnant rodents fail to form a decidua, indicating that a focal object alone is insufficient (Alden and Smith, 1959). In 1963, it was determined that large volumes of oils (10μl-100μl) can induce decidualization (Finn and Keen, 1963; Wang et al., 2020) and this finding was perplexing as oil did not represent a focal object such as an embryo or a bead. However, if the volume of oil is reduced to 1μl focal implantation sites can be detected (Chen et al., 2011). Later it was also proposed that lectins (proteins that can bind to carbohydrates) including Concanavilin A (ConA), Wheat Germ Agglutinin (WGA) and Soybean lectin, ConA coated beads and calcium ionophore A23187 can induce formation of decidua (Buxton and Murdoch, 1982; Herington et al., 2009; Sakoff and Murdoch, 1994). More recently, it has been proposed that a threshold amount of air when injected into the uterine lumen through the cervix can also induce decidua formation (Li et al., 2019). Thus, a variety of artificial stimuli are capable of inducing decidualization during the receptive period.

Ovarian signals are key to uterine receptivity across mammalian species. In mice, these signals have been well-studied using an ovariectomy model with a hormone regimen mimicking estrous cycle hormones (Peterse et al., 2018). This model shows that implantation and subsequent decidualization absolutely rely on a small estrogen surge before implantation. It has also been used to test various artificial decidualization models. While most stimuli require both progesterone and some estrogen for the formation of a decidua, the trauma stimulus (using a needle to scratch the anti-mesometrial pole of the uterine lumen) can induce decidualization with progesterone alone (Finn, 1966). Thus, of all the models, trauma induced decidualization is thought to be the least representative of the embryo-induced decidualization. In addition to the ovarian hormones, all models of artificial decidualization require physical contact between the luminal epithelium and the stroma prior to application of stimulus for decidua formation (Lejeune et al., 1981). If such contact is severed enzymatically or mechanically, decidualization does not occur even with a trauma stimulus supporting the need for a physical connection between the epithelial and stromal cell types (Lejeune et al., 1981).

Evidence for pathways necessary for decidualization comes from experiments where administration of a substance during the receptive period interferes with implantation and decidualization. Treatment with tranylcypromine – a prostacyclin inhibitor, blocks artificial decidualization (Buxton and Murdoch, 1982) and this is attributed to loss of prostacyclins, catecholamines and luteolytic prostaglandins. Further, artificial decidualization can be blocked using indomethacin and propranolol, suggesting that prostaglandins and uterine beta-adrenergic receptors are critical for decidualization. Other studies also suggest a critical role for prostaglandins in decidualization (Rankin et al., 1981; Scherle et al., 2000) and PTGS2 is expressed in the sub-epithelial stroma under the mouse implantation chamber and is critical for embryo growth and continued decidualization (Massri and Arora, 2025). The polyphosphatidylinositol pathway and calcium are also implicated in the induction of the decidua (Kyd and Murdoch, 1992; Sakoff and Murdoch, 1994). Further, intraluminal administration of ConA can inhibit implantation suggesting that the embryo must recognize certain sugar molecules on the endometrial surface in order to implant (Hicks and Guzmá-González, 1979). In support of this, the N-Acetyl Glucosamine group is expressed in the uterine epithelium during the peri-implantation period (Munson et al., 1989; Yasunaga et al., 2012).

When comparing artificial models against embryo-mediated decidualization, the ConA-bead model and oil model both expressed TJP1, cadherin 1 (CDH1), and CTNNB1 in the luminal epithelium of the decidualized uterus (Herington et al., 2009). Further, an avascular zone near the ConA-bead attachment site was observed similar to embryo-mediated decidualization (Herington et al., 2009). Differential gene expression analysis (McConaha et al., 2011) between bead implantation site and inter-implantation site showed upregulation of key implantation related transcripts including *Akp2, Bmp2, Bmp8a, Wnt4, Fkbp5, Cebpb and Ptgs2* (Herington et al., 2009). Thus, of the available models of decidualization ConA-bead model of artificial decidualization is thought to more closely resemble the embryo’s stimulus. These data also suggest that there is a connection between site of embryo/bead attachment and induction of decidua formation.

With respect to the formation of an implantation chamber, it has been shown that while ConA-beads or agarose beads coated with HBEGF can induce formation of an implantation chamber agarose beads alone cannot (Sakurai et al., 2024; Yuan et al., 2018). This suggests that a physical stimulus with certain biochemical properties can initiate both chamber formation and decidualization. However, the relationship between implantation chamber formation and the onset of decidualization remains unclear. One of the oldest evidence connecting implantation to decidualization was provided by Finn and Mclaren, where they suggested that blue dye implantation sites appear first, followed by alkaline phosphatase positive stroma – suggesting that implantation (detected by blue dye sites) precedes decidualization (detected by stromal alkaline phosphatase expression – indicator of epithelialization) (Finn and McLaren, 1967). In a separate study, it was shown that coating agarose beads with HBEGF and IGF1 induces vascular permeability, decidualization and expression of Bmp2 at the sites of bead attachment (Paria et al., 2001) connecting the site of physical stimulus with implantation and decidualization.

Previous research from our lab evaluating early events in murine pregnancy suggests that once the embryos enter the uterus, they move to the center of the uterine horn (Flores et al., 2020). At this time, the uterine tube displays dynamic folding patterns with primarily longitudinal folds that run along the oviductal-cervical axis (Madhavan et al., 2022). Longitudinal folds transition into transverse folds along the mesometrial-anti-mesometrial axis when the embryos are located in the middle of the horn. Scattering of embryos from the center towards the oviduct and the cervical end of the uterus induces formation of flat peri-implantation regions (PIRs). The center of these peri-implantation regions is the implantation site where an embryo will eventually attach and initiate formation of an implantation chamber followed by decidualization of stromal cells around the chamber (Madhavan et al., 2022; Massri and Arora, 2025). While these data suggest a sequence of events in which the implantation chamber precedes decidualization, it is unknown whether chamber formation is a prerequisite for inducing decidualization. In species such as humans and bats decidualization initiates independent of the embryo (Jarrell, 2018). Thus, while reporting the impact of genetic mutations on pregnancy, researchers first perform tests for embryo implantation using the tail vein blue dye injection. Irrespective of the data obtained researchers conduct a test for decidualization by introducing oil in a pseudopregnant uterus (Wang et al., 2020). In this case, if decidualization fails the phenotype is attributed to defects in stromal cell decidualization. However, if implantation chamber precedes decidualization, implantation failure will always be followed by decidualization failure.

To address whether chamber formation is critical for initiating decidualization in the mouse and to determine what aspects of implantation and decidualization are dependent on a physical stimulus vs molecular dialogue we turned to commonly used models of artificial decidualization. These include oil and ConA-beads that induce decidualization and models that fail to form a decidua – agarose blue beads and eggs. We observed that stimuli that induce artificial decidualization initially form a V-shaped implantation chamber with PTGS2 expression in the subepithelial stroma under the chamber. As decidualization progresses, PTGS2 expression shifts from beneath the chamber to laterally around the chamber. We also demonstrate that other than ConA, beads coated with lectins that share the sugar binding properties of ConA can also induce a V-shaped chamber. Finally, we show that irrespective of the physical stimulus that induces the implantation chamber, collagen fibers in the decidualizing stroma are oriented radially away from the chamber. Our data conclusively suggest that focal stimuli mediated implantation chamber formation precedes stromal cell decidualization.

## RESULTS AND DISCUSSION

### Pre-implantation uterine folding patterns are independent of the focal stimulus

Remodeling of the uterine epithelium into pre-implantation transverse folds is critical to prevent embryos from being trapped in longitudinal folds at the time of implantation (Madhavan et al., 2022). To determine if an embryo as a molecular signal is necessary for transverse folding, we examined uterine folding patterns at GD3.5 with various stimuli. In naturally mated mice we observed transverse epithelial folds along the mesometrial-anti mesometrial axis (**Fig. 1A, A’**) and found embryos clustered in the center of the lumen (**Fig. 1A’’**) (Flores et al., 2020). To assess the relationship among epithelial folding, embryo number, and uterine horn length, we calculated the coefficient of determination (**Fig. 1B-D**). We observed that while the length of the uterine horn correlates highly with both the number of folds (R^2^=0.8215) and the number of embryos (R^2^=0.7817), the correlation between the number of embryos and the number of folds is low (R^2^=0.3697). This suggests that embryo number does not directly affect the fold number. The correlation between the uterine horn length and the embryo number may be explained by levels of circulating Estrogen (E2) at the onset of pregnancy. The levels of E2 depend on the number of ovulated follicles (Schütz and Batalha, 2024). A higher number of ovulated eggs may thus increase E2 levels, leading to higher E2-Estrogen Receptor 1 (ESR1)–dependent proliferation of epithelium, muscle, and elongation of uterine horn. Intuitively, a longer uterine tube must be partitioned into a higher number of folds and PIRs, however, the underlying mechanisms of uterine fold formation are less well understood and should be a subject of future investigations.

**Figure 1:**
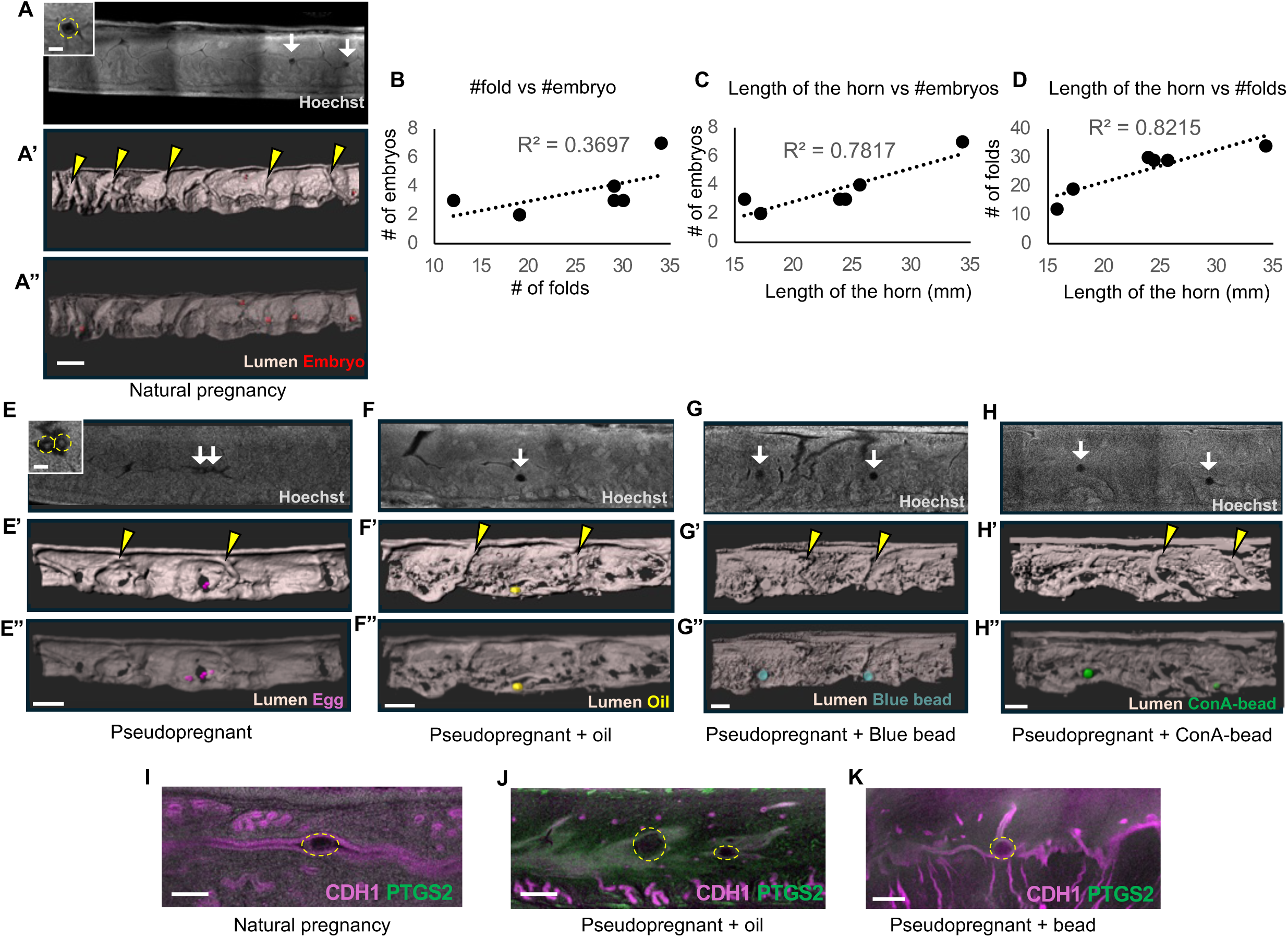
Preimplantation uterine epithelial folding occurs independent of object type. **(A-A’’)** Optical slice from confocal imaging at GD3.5 of natural pregnancy (n=4 mice) with embryos **(A)** and 3D reconstruction of the lumen with transverse folds **(A’)** and embryo surfaces (in red) **(A”).** Inset in **A** shows an embryo. **(B-D)** Coefficient of determination between #folds vs #embryo **(B)**, length of the horn vs #embryos **(C)**, length of the horn vs #folds **(D)** (n=6 mice, 6 uterine horns), black dotted line indicates the linear trendline. **(E-H)** Optical slice from confocal imaging **(E, F, G, H)** and 3D reconstruction of the lumen **(E’, F’, G’, H’)** of a pseudopregnant uterus with eggs (in pink) (n=3 mice, n_o_=17) **(E”)**, oil (in yellow) (n=3 mice, n_o_=20) **(F”)**, blue beads (in blue) (n=4 mice, n_o_=9) **(G”),** and ConA-beads (in green) (n=4 mice, n_o_=9) **(H”)**. Inset in **E** shows eggs. Yellow arrowheads: transverse folds, white arrows: object location. **(I-K)** No PTGS2 expression observed in uterus containing an embryo (n=3 mice, n_o_=10) **(I)** or a bead (n=4 mice, n_o_=7 per bead type) **(K)**. PTGS2 expression observed in the lumen when oil (n=4 mice, n_o_=9) **(J)** was transferred. Yellow dotted circles: embryo/objects. n_o_-number of objects. Scale bars: **A,A’,A”:** 400 µm, **A,E inset:** 100 µm, **E,E’,E”,F,F’,F”,G,G’,G”,H,H’,H”:** 300 µm, **I:** 150 µm, **J:** 200 µm, **K:** 250µm.

In pseudopregnant uteri, where no embryos were present, we also observed transverse epithelial folds throughout the length of the uterus at GD3.5 (**Fig. 1E, E’**). Concurrent with previous observations we detected eggs in the pseudopregnant uteri (McLaren, 1970) (**Fig. 1E”**). When pseudopregnant uteri were stimulated with oil and beads, we observed spherical empty spaces in the case of oil (presumed to be oil droplets), (**Fig. 1F**) and beads (**Fig. 1G, H**) in uteri. Intriguingly, when oil or beads were transferred to the pseudopregnant uteri, eggs were never detected. This observation resembles cases where embryos are detected in the uterine horn while eggs remain undetected, likely due to differences in the timing of embryo versus egg entry into the uterus (McLaren, 1970). All uteri where objects were transferred showed transverse epithelial folds at GD3.5 (**Fig. 1F’, G’, H’**). These data suggest that uterine transverse folding is an autonomous process likely dependent on ovarian function and uterine contractions but independent of the type of object present in the uterine horn. The lumen around the oil droplets or beads was closed whereas eggs were seen free-floating in the uterine lumen. This luminal closure may be a function of the object size, with eggs being <80μm (Alberts et al., 2002) and blastocysts or beads are >100μm (Levron et al., 1996). Alternatively the zona pellucida on the egg may restrict contact with the luminal epithelium thereby preventing luminal closure (Alberts et al., 2002).

To determine the response of the luminal epithelium to the type of object, we stained for PTGS2 (**Fig. 1I-K**). Consistent with previous findings (Massri and Arora, 2025) we observed PTGS2 staining in the luminal epithelium throughout the uterine horn when oil was injected (**Fig. 1J**). However, in the presence of embryos (**Fig. 1I**) or any kind of bead (**Fig. 1K**), PTGS2 expression was not observed in the uterus. These data suggest that oil has certain properties, either lipid-like nature or sulphated polysaccharides, to induce PTGS2 expression in the lumen (Massri and Arora, 2025). Expression of such high levels of PTGS2 may also preemptively cause prostaglandin secretion which should be considered when using oil as a model for implantation and decidualization studies.

### Concanavalin A beads and oil droplets induce implantation chamber formation

Humans can initiate predecidualization without the presence of an embryo; however, complete decidualization requires embryonic signals (Okada et al., 2018). In contrast, rodents need either an embryo or an artificial stimulus to trigger decidualization. While objects or oil can induce a decidual response in rodents that resembles natural pregnancy (Herington et al., 2009), it remains unclear whether these artificial stimuli also lead to the formation of implantation chambers. To determine which type of objects can trigger formation of an implantation chamber we used different models of pregnancy. In naturally mated mice, once the embryo arrives at the implantation site, a V-shaped implantation chamber begins to form (Madhavan et al., 2022) (**Fig. 2A’**). Implantation chambers are flanked by inter-implantation sites that continue to show transverse epithelial folds (Madhavan et al., 2022). Implantation chamber formation is also coincidental with increase in vascular permeability at the implantation site as observed by intravenous blue dye injections (**Fig. 2A**). At GD4.5 pseudopregnant uteri no longer showed the presence of eggs, suggesting that eggs likely degrade prior to GD4.5. Further no implantation chambers were observed in pseudopregnant uteri at GD4.5 (**Fig. 2B**). Unlike the organized distribution of flat and folded regions in the pregnant uterus, pseudopregnant uteri exhibited either transverse folds alone or folds in transition (**Fig. 2B’**). These data suggest that in the absence of an implantation event the uterine lumen begins to transition away from its transverse-only fold pattern likely in preparation for the next cycle. Pseudopregnant uteri injected with blue beads failed to show blue dye implantation sites (**Fig. 2C**). These uteri continued to show the presence of the beads at GD4.5 however no implantation chambers were observed (**Fig. 2C’**). These data suggest that blue beads can make contact with the lumen however there is absence of a molecular dialogue between the bead and the epithelium that prevents implantation chamber initiation.

**Figure 2:**
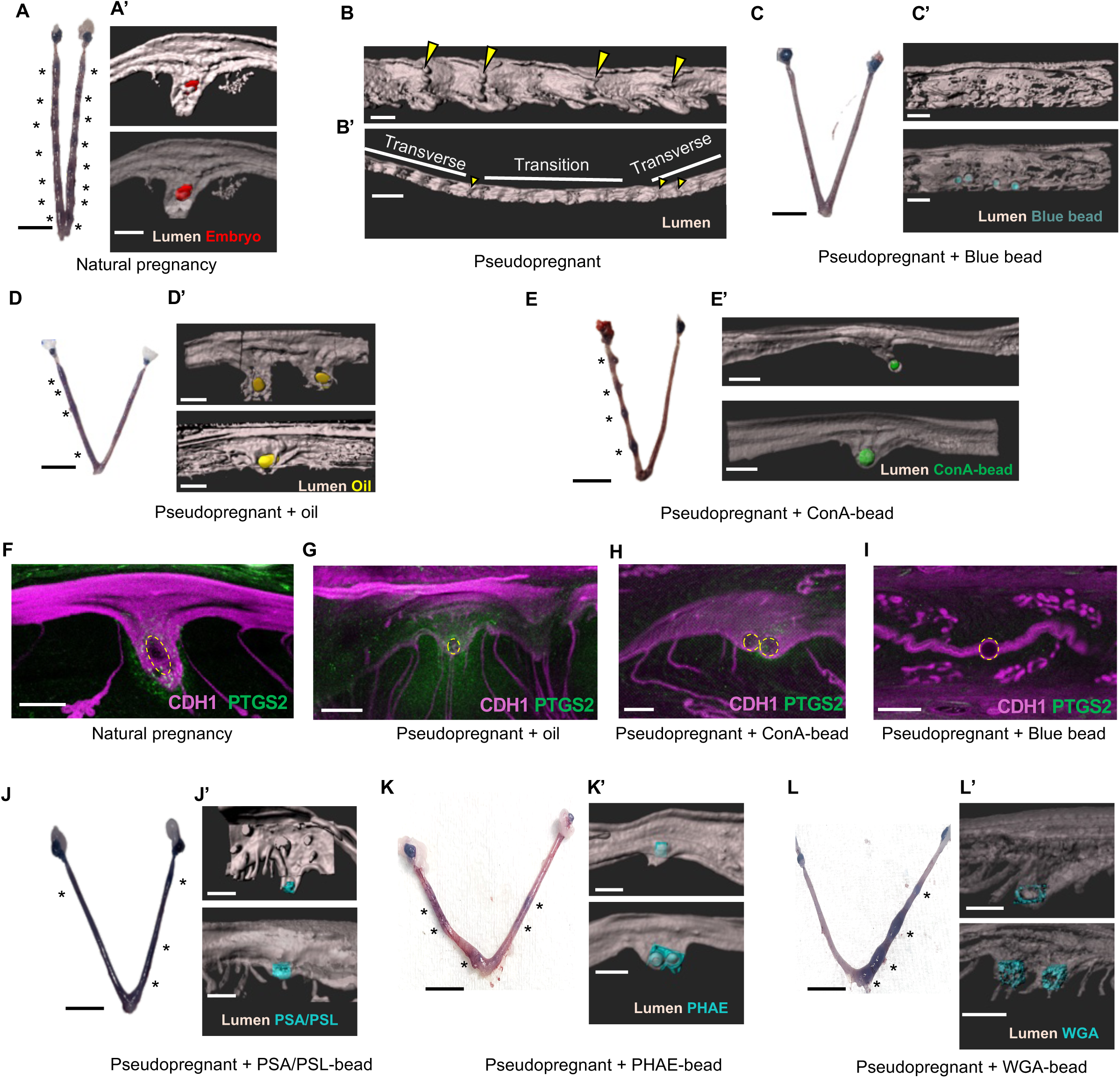
Embryo, oil, and ConA-beads can initiate implantation chamber formation. **(A-A’)** At GD4.5, implantation in mice of natural pregnancy was confirmed with blue dye sites **(A)**. 3D reconstruction of the implantation site where embryos (in red) form a V-shaped chamber **(A’)** (n=4 mice, n_o_=14). **(B-B’)** 3D reconstructed lumen of a pseudopregnant mouse with transverse folds (indicated with yellow arrowheads) **(B)** and a partial region of folds that are transitioning away from transverse folds (n=4 mice) **(B’)**. **(C-C’)** No blue dye reaction sites was observed when blue beads were transferred into pseudopregnant mice **(C).** 3D reconstruction shows no implantation chamber formation with blue beads (in blue) (n=3 mice, n_o_=10) **(C’)**. **(D-E)** Blue dye sites were identified when oil (n=3 mice) **(D)** and ConA-beads (n=4 mice) **(E)** were transferred into pseudopregnant uteri. **(D’-E’)** 3D reconstruction of the implantation site with oil (in yellow) (n=3 mice, n_o_=9) **(D’)** and ConA-beads (in green) (n=4 mice, n_o_=14) **(E’)**. **(F-I)** PTGS2 expression observed in stroma under the implantation chamber with embryo (n=2 mice, n_o_=6) **(F),** oil (n=4 mice, n_o_=9) **(G),** and ConA-beads (n=2 mice, n_o_=9) **(H)**, but not with blue beads (n=3 mice, n_o_=6) **(I)**. Yellow dotted circles: embryo/objects. **(J-L)** Blue dye sites were identified when lectin-coated beads were transferred in pseudopregnant uteri. PSA/PSL beads (n=3 mice) **(J)**, PHAE beads (n=3 mice) **(K)** and WGA beads (n=3 mice) **(L). (J’-L’)** 3D reconstruction of the implantation chambers induced with lectin-coated beads (in cyan). PSA/PSL (n=3 mice, n_o_=3) **(J’)**, PHAE (n=3 mice, n_0_=4) **(K’)** and WGA (n=3 mice, n_o_=5) **(L)**. Asterisk indicates implantation site. n_o_-number of objects. Scale bars: **A’, B, D’, I:** 200 µm, **B’:** 1000 µm, **C’, E’, F:** 300 µm, **G:** 500 µm, **H:** 150 µm, **A,C,D,E,J,K,L:** 5mm.

Similar to naturally mated mice with embryos, pseudopregnant uteri that were injected with low volume oil (**Fig. 2D**) and ConA-beads (**Fig. 2E**) showed blue dye implantation sites. When 3D surfaces were constructed, they showed formation of V-shaped implantation chambers at the site of the object (**Fig. 2D’, E’**). Although it has been previously shown that ConA-beads can form implantation chambers those beads were injected at GD3.5 (Sakurai et al., 2024) while embryos typically enter the uterus at the morning of GD3. We observed that not all ConA-beads or oil droplets induced formation of an implantation chamber (**Supp Fig. 2)**. Uneven spacing of oil droplets and ConA-beads was observed, in contrast to the uniform embryo distribution seen in natural mating (**Supp Fig. 2)** (Flores et al., 2020). These data suggest that communication between embryos and/or communication between the embryo and the uterus is critical for even spacing of objects throughout the uterine horn. However, an implantation chamber can be induced by both ConA-beads and oil droplets. We have previously shown that elongation of the implantation chamber is necessary for the embryo to align its embryonic-abembryonic axis along the uterine mesometrial-anti-mesometrial axis (Madhavan et al., 2022). Since oil and ConA-bead can both form an implantation chamber (**Fig. 2D’, E’**) and neither has an axis or polarity, these findings further suggest that implantation chamber elongation precedes embryo rotation and axis alignment in naturally mated embryos (Madhavan et al., 2022).

In an embryo-induced implantation chamber, PTGS2 expression is observed in the stroma underlying the chamber (Massri and Arora, 2025) (**Fig. 2F**). We observed similar PTGS2 expression near the chamber in oil treated (**Fig. 2G**) and ConA-bead treated uteri (**Fig. 2H**). However, PTGS2 expression was absent in uteri injected with blue beads (**Fig. 2I**). These data confirm that stromal PTGS2 expression is an indication of successful crosstalk between the object and the luminal epithelium to initiate changes in the underlying stroma in the context of an implantation chamber. Further, in the absence of an implantation chamber, PTGS2 expression (and initiation of decidualization) is not observed.

For oil, sulphated polysaccharides in oil are proposed as the stimulus necessary for inducing decidualization and both agar and heparin that have sulphated polysaccharides can also induce formation of a decidua (Finn and Keen, 1963). For the beads, since ConA-beads are able to induce a chamber but blue sepharose beads are not, we hypothesized that lectin-like properties of ConA may contribute to its ability to act as an inducer of implantation chamber formation and initiator of decidualization. ConA is characterized to have a sugar specificity for branched N-linked oligosaccharides, N-acetyl glucosamine and mannose (Cummings and Etzler, 2009). We identified other lectins that may share some of these properties. Wheat Germ Agglutinin (WGA) has a sugar specificity for N-acetyl glucosamine, *Pisum Sativum* Agglutinin (PSA) binds to a subset of N-linked oligosaccharides and mannose, and *Phaseolus vulgaris* erythroagglutinin (PHA-E) is specific for biantennary galactosylated N-glycans containing a bisecting N-acetyl glucosamine residue (Cummings and Etzler, 2009). We observed that all 3 lectins WGA, PSA and PHA-E were able to induce blue dye implantation sites (**Fig. 2J, K, L**) and V-shaped implantation chambers at GD4.5 (**Fig. 2J’, K’, L’**). These data suggest that uterine luminal cells express sugars that are recognized by the different lectins and likely the trophoblast cells of the embryo. Epithelial sugar binding to the object may facilitate formation of an implantation chamber and induction of stromal PTGS2.

### Implantation chamber elongation and decidualization observed with Concanavalin A beads and oil droplets in the uterus

To ascertain the connection between implantation chamber growth and decidualization we evaluated pregnancy progression in the naturally mated uteri and pseudopregnant uteri injected with oil and ConA-beads. At GD5.5 when decidual sites are easily visible in the naturally mated uteri (**Fig. 3A**) we observed that the chambers continue to elongate (Massri and Arora, 2025) (**Fig. 3A’**). We observed similar decidual sites (**Fig. 3B, C)** and chamber elongation in pseudopregnant uteri injected with oil (**Fig. 3B’**) or ConA-beads (**Fig. 3C’**). PTGS2 expression in a natural mating scenario was observed on the lateral sides of the implantation chamber and at the mesometrial pole of the chamber (**Fig. 3D**). PTGS2 expression with the oil droplet seemed to mirror that observed with the embryo (**Fig. 3E**) however with ConA-beads, the PTGS2 expression was largely in the stroma at the anti-mesometrial pole of the chamber (**Fig. 3F**). These differences might be due to the flexibility of oil droplets to change shape and elongate along with the chamber similar to embryo elongation to an epiblast at this stage (Massri and Arora, 2025). ConA-beads on the other hand are rigid and unable to elongate along with the chamber. Since embryos are evenly spaced but beads and oil drops do not space out evenly, multiple implantation chambers with ConA-beads (**Fig. 3C’**) were frequently observed. To study chamber elongation and resulting PTGS2-mediated decidualization, low volume oil might be a better model compared to other models of artificial decidualization.

**Figure 3:**
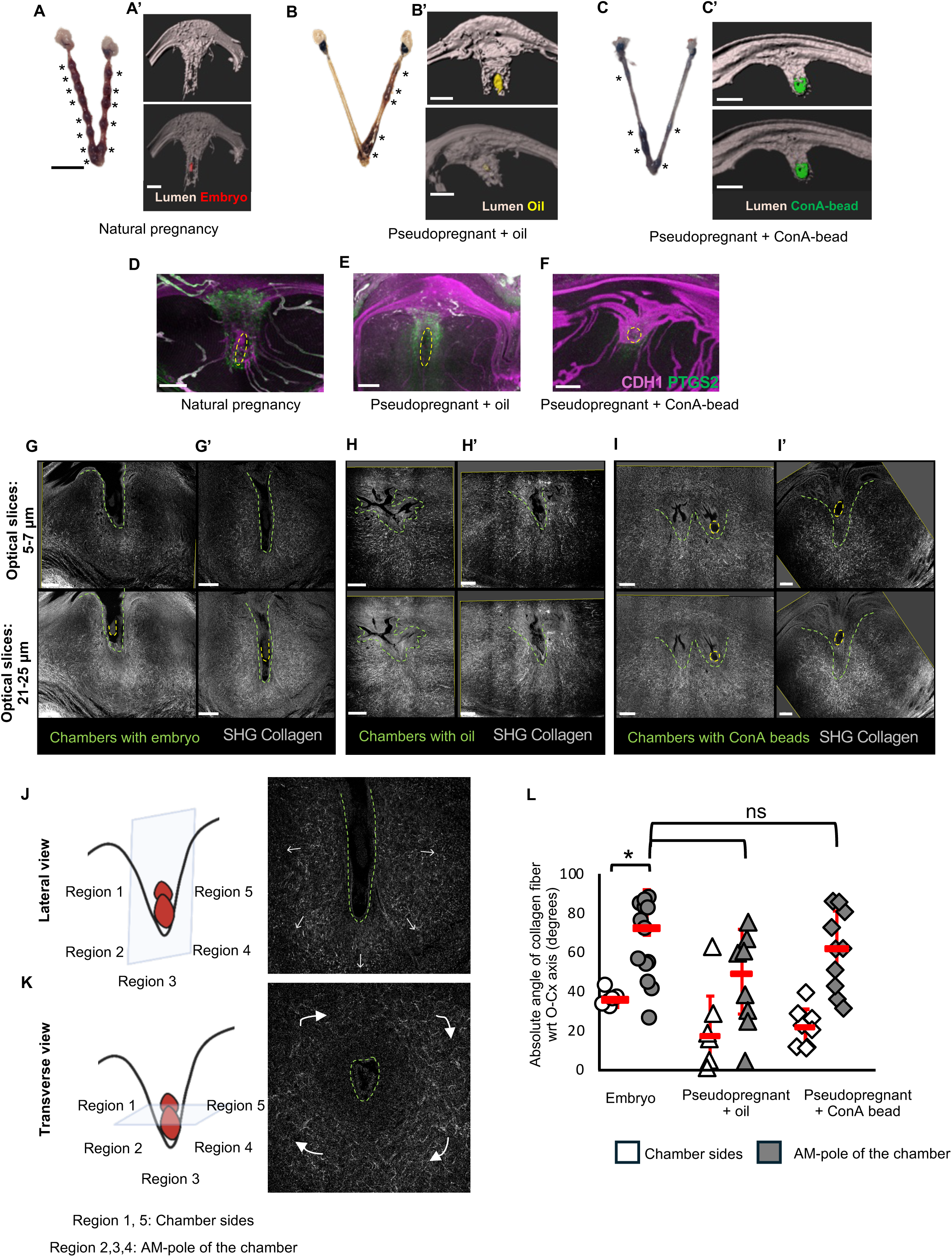
Embryo, oil, and ConA-beads induce elongated implantation chambers and decidualization at GD5.5. **(A-C)** Decidual sites were observed with blue dye for embryos (n=5 mice) **(A)**, oil (n=5 mice) **(B)**, and ConA-beads (n=3 mice) **(C)**. Asterisk indicates decidual site. **(A’-C’)** 3D reconstruction of the V-shaped chamber with embryos (in red) (n=5 mice, n_o_=9) **(A’)**, oil (in yellow) (n=5 mice, n_o_=9) **(B’)** and ConA-bead (in green) (n=3 mice, n_o_=11) **(C’)**. **(D-F)** Comparable PTGS2 expression observed in decidualizing stroma with embryo (n=4 mice, n_o_=5) **(D)** and oil (n=4 mice, n_o_=7) **(E)**. Implantation chamber with ConA-bead (n=3 mice, n_o_=11) **(F)** shows PTGS2 expression only in the stroma at the AM pole of the implantation chamber. Yellow dotted circles indicate embryo/objects. **(G-I’)** Optical slice views of the SHG signals (grey) from multiphoton imaging of collagen fibers in stroma surrounding implantation chambers at GD5.5 induced by embryo (n=3 mice) **(G,G’)**, oil (n=4 mice) **(H,H’)** and ConA-beads (n=3 mice) **(I, I’)**. Green dotted line: chamber, yellow dotted line: embryo/object. **(J-K)** Orientation of collagen fibers depends on the plane of sectioning. Collagen fibers orient along the invading embryos when viewed laterally **(J)** and wrapped around the chamber when viewed transversely **(I)**. White arrows indicate the direction of fiber orientation. **(L)** Quantification of collagen fiber orientation angle with uterine oviductal-cervical axis for embryo (n=3 mice, n_c_=4), oil (n=4 mice, n_c_=9) and ConA-beads(n=3 mice, n_c_=7) (\**p*<0.1, Wilcoxon Signed-Rank Test for comparing within embryo; Mann-Whitney U test for comparing embryo vs oil/ConA-bead). Red dashes indicate the median angle. n_o_-number of objects, n_c_-number of chambers, SHG – Second Harmonic Generation. Scale bars: **A’,B’,C’:** 200 µm, **D:** 150 µm, **E,F:** 200 µm, **G,G’,H,H’:** 200 µm, **I, I’:** 100µm, **A,B,C:** 5mm.

Extracellular matrix especially collagen organization has been reported to be distinct based on ovarian hormones, pregnancy and the particular stage of pregnancy (Fainstat, 1963). More recently, Second Harmonic Generation (SHG) has been used for visualization of fibrillar collagen types I, III, and V in the decidualizing stroma around the implantation chamber (Savolainen et al., 2024). It has been proposed that SHG signals are only visible in the secondary decidual zone and oil-mediated artificial decidualization results in disorganized fibrils compared to organized fibrils in embryo-mediated decidualization (Savolainen et al., 2024). Another study suggests that SHG signals are detected even prior to embryo implantation, however the organization of the collagen fibers as observed by SHG is different in response to the implantation chamber and decidualization (Gebril et al., 2025). We used SHG to ascertain collagen fibril organization around the object-mediated implantation chamber. We determined that it was critical to compare fibril organization in natural and artificial models of decidualization by keeping the orientation of the chamber comparable. When sliced along the M–AM axis, collagen fibrils were organized radially outward from the V-shaped implantation chamber, irrespective of the object (embryo, ConA-bead, or oil) inducing the chamber (**Fig. 3G, G’, H’, I, I’**). In the case of oil-mediated decidualization, irregular chamber formation that did not represent distinct V-shaped chambers was accompanied by irregular collagen fibril organization (**Fig. 3H**). When quantified we observed that fibrils around the chamber sides had smaller angles with respect to the oviductal-cervical axis and fibrils near the AM pole stroma had larger angles irrespective of the object used to induce the chamber (**Fig. 3L**) These data suggest that collagen fibril organization is a function of chamber shape. Depending on the plane of the optical section around the embryo fibril orientation appears distinct (**Fig. 3J, K**). Further, it is possible that the fibrils start out disorganized as is the case with wound healing (Stroncek and Reichert, 2008) but they quickly organize in the radial orientation to support formation of the expanding decidua.

### Implantation chamber formation precedes decidualization

Oil, ConA-beads and other lectin-coated beads as focal physical stimuli are capable of inducing implantation chamber formation and decidualization in a hormone primed uterus. We determined that uterine folding is independent of the type of object present in the uterus. Chamber initiation and communication between the epithelium and the stroma is dependent on sugar-like properties of lectins and/or lipid like nature of oils. The ability of lectins to regulate object detection by the uterine lumen may be a conserved mechanism for species-specific embryo-uterine interactions (Wang et al., 2023) and can be explored in future studies. An important concept that emerges out of studying these various models of decidualization is that as is the case with the embryo, implantation chamber formation is necessary to initiate epithelial-mesenchymal communication and to induce decidualization. These observations are specific to the E2-ESR1 signaling dependent models of decidualization as we did not test the scratch model of decidualization that has been shown to be independent of E2-ESR1 signaling. A better understanding of the physical stimuli-dependent, embryo-dependent and embryo-independent stimuli that are critical for initiation of decidualization can give way to novel therapeutic approaches for treating clinical conditions like recurrent implantation failure and also for developing novel contraceptives.

## ACKNOWLEDGEMENTS

We thank Sara Makaremi for help with the multiphoton microscope for SHG imaging and Manoj Madhavan for critical review of the manuscript.

## AUTHOR CONTRIBUTIONS

HRK and RA conceptualized the study and designed the experiments. HRK, NM, AVB, AK, PG performed the experiments. HRK, NM, AK and RA validated the data and performed the analyses. HRK, AVB and RA prepared the figures and wrote and edited the manuscript. All authors reviewed and accepted the final version of the manuscript.

## GRANT FUNDING

This research was supported in part by the NIH R01HD109152 and March of Dimes grant #5-FY20-209 to R.A., the Eunice Kennedy Shriver National Institute of Child Health & Human Development of the National Institutes of Health under award #T32HD087166 to H.R.K and N.M., and the Walstrom Family Endowed Women’s Health Research Award by Dept. of Ob/Gyn and Reproductive Biology at Michigan State University to A.V.B.

## CONFLICT OF INTEREST STATEMENT

The authors declare no conflict of interest

## METHODS

### Mouse models of natural and artificial pregnancy and decidualization

6-10 week old CD1 wild-type mice were used for these studies. For pregnancy studies, females were mated with fertile males (**Supp Fig. 1A**). To induce pseudopregnancy, females were mated with vasectomized males (**Supp Fig. 1B**). The appearance of a vaginal plug was identified as a gestational day (GD) 0.5 (GD0 1200h). To induce artificial decidualization, blue beads (#153-7302, Bio-Rad), Concanavalin A (ConA) beads (C9017, Sigma), or 1 µl sesame oil (241002500, Thermo Scientific) with 3 µl PBS were injected into the uterine lumen either surgically or using a non-surgical embryo transfer (NSET) device (60020, mC&I Device for Mice) into a pseudo-pregnant mouse at GD2 1800h (**Supp Fig. 1C**). Uteri were dissected from mice at GD3.5 (GD3 1200h-1400h), GD4.5 (GD4 1200h – 1800h) and GD5.5 (GD5 1200h-1400h). To assess the role of lectin beads coated with Wheat Germ Agglutinin (WGA) (Bioworld 20181018-5), *Pisum Sativum* Agglutinin (PSA) (Bioworld, 21510408-2) and *Phaseolus vulgaris* erythroagglutinin (PHA-E) (Bioworld, 30330025-1) were injected into the mice at GD2 1800h and uteri were dissected at GD4.5. To detect implantation sites, a 0.25 ml intravenous injection of 1.5% Evans blue dye (MP Biomedicals, ICN15110805) was administered 10 minutes before euthanizing mice at GD4.5 and GD5.5. All mice were maintained on a 12-hour light/dark cycle, and all mouse studies and protocols were approved by the Institutional Animal Care and Use Committee at Michigan State University.

### Whole-mount immunofluorescence staining

After dissections, the uteri were fixed in a 1:4 solution of DMSO: Methanol and stored at −20 °C. Whole mount staining of the entire uterine horns was performed as described previously (Arora et al., 2016; Flores et al., 2020; Madhavan et al., 2022). Briefly, samples were rehydrated in a (1:1) methanol: PBST (PBS, 1% Triton (Sigma Aldrich, T9284)) solution for 15 minutes, followed by a 15 minutes wash in PBST. Samples were then incubated in a blocking solution (PBS, 1% Triton, and 2% powdered milk) for 1 hour at room temperature, followed by incubation with primary antibodies in the blocking solution for seven nights at 4°C. Primary antibodies used in this study include Rat anti-CDH1 (Takada, M108, 1:500), Rabbit anti-FOXA2 (Abcam, ab108422, 1:500) and Rabbit anti-PTGS2 (Abcam, ab109025, 1:200). Uteri were then washed in PBST solution for 2X15 minutes and 4X45 minutes and incubated with Hoechst and Alexa Fluor-conjugated secondary antibodies (1:500) for three nights at 4°C. Secondary antibodies used were Donkey anti-Rabbit 555 (A31572, Invitrogen) and Goat anti-Rat 633 (A21094, Invitrogen). Uteri were then washed with PBST for 1X15 minutes and 3X45 minutes, followed by dehydration in methanol for 15 min. Uteri were then bleached in a 3% H_2_O_2_ solution (Sigma-Aldrich 216763) prepared in methanol overnight at 4°C. The samples were washed with 100% methanol for 3X30 minutes and cleared in a 1:2 solution of benzyl alcohol: benzyl benzoate (1:2) (Sigma-Aldrich, 108006, B6630) overnight.

### Confocal Microscopy

All samples were imaged on a Leica SP8 TCS white light laser confocal microscope with a 10x air objective and 7.0 mm Z stack using Leica Application Suite X (LAS X) software (version 3.5.7.23225) (**Supp Fig. 1D**).

### Image segmentation to visualize uterine lumen, implantation chamber, embryos and objects

Upon imaging, the confocal image files (.LIF format) were imported into the 3D surpass mode Imaris v9.2.1 (Bitplane; Oxford Instruments, Abingdon, UK). To visualize the uterine lumen and the implantation chamber the FOXA2+ gland epithelium signal was subtracted from the total epithelial CDH1 fluorescent signal. The Hoechst signal was used to locate eggs or embryos. 3D renderings were created using the SURFACE module as described previously (Madhavan et al 2022). Snapshots were taken with variable degree of transparency for the lumen surface in order to highlight the objects as shown in the figures. (**Supp Fig. 1 E,F**)

### Quantifying fold numbers

The entire lumen was reconstructed using the SURFACE module as previously described(Arora et al., 2016). To identify highly folded regions, a MATLAB script for surface curvature analysis(Arora et al., 2016) was applied to the lumen surfaces. Folded regions were manually counted for each uterine horn. Embryos were identified with Hoechst staining and counted manually.

### Second Harmonic Generation (SHG)

Whole uteri stored in benzyl alcohol: benzyl benzoate were utilized in this experiment. A Leica SP8 DIVE laser multiphoton microscope equipped with Spectra-Physics Insight X3 dual beam (680 to 1,300 nm tunable and 1,040 nm fixed) and 4Tune, tunable, super sensitive hybrid detectors (HyDs) with a 25x objective lens was employed to acquire images of the decidual site (LAS X software version 3.5.7.23225). A wavelength of 950 nm was used to detect a collagen signal at 475 nm. The acquired images were subsequently analyzed using Imaris v9.2.1. (Bitplane; Oxford Instruments, Abingdon, UK).

### Quantification of collagen fiber orientation angle from SHG images

Following SHG imaging, 5-7mm images of the decidual site in the M-AM plane was captured using the snapshot function in Imaris v9.2.1. The .tiff image was loaded into ImageJ (v2.16.0/1.54p). For fiber orientation, the image was first converted to 8-bit and then analyzed for local orientation of fiber angles with respect to the implantation chamber using the OrientationJ plugin. The Orientation J Analysis plugin was used for color coding the fibrils and the OrientationJ Measure plugin was used for measuring the angle of collagen fibers for selected 4-5 regions of interests (ROIs) around the implantation chamber. Absolute angles for regions on the left/right of the chamber or at the anti-mesometrial (AM) pole of the chamber were pooled and plotted for different decidualizing signals (**Supp. Fig. 3**).

### Statistics

Statistical analysis was performed using Microsoft Excel and GraphPad Prism v10. The coefficient of determination (R²) was calculated from Pearson’s correlation coefficient to test pairwise correlations between the number of folds, number of embryos, and length of the lumen. Wilcoxon Signed-Rank Test for comparing SHG data fiber orientation between left/right and AM pole data for the embryo chamber and Mann-Whitney U test was used for comparing SHG data of embryo with oil and ConA-bead induced chambers.

---

**Supplementary Figure 1.**
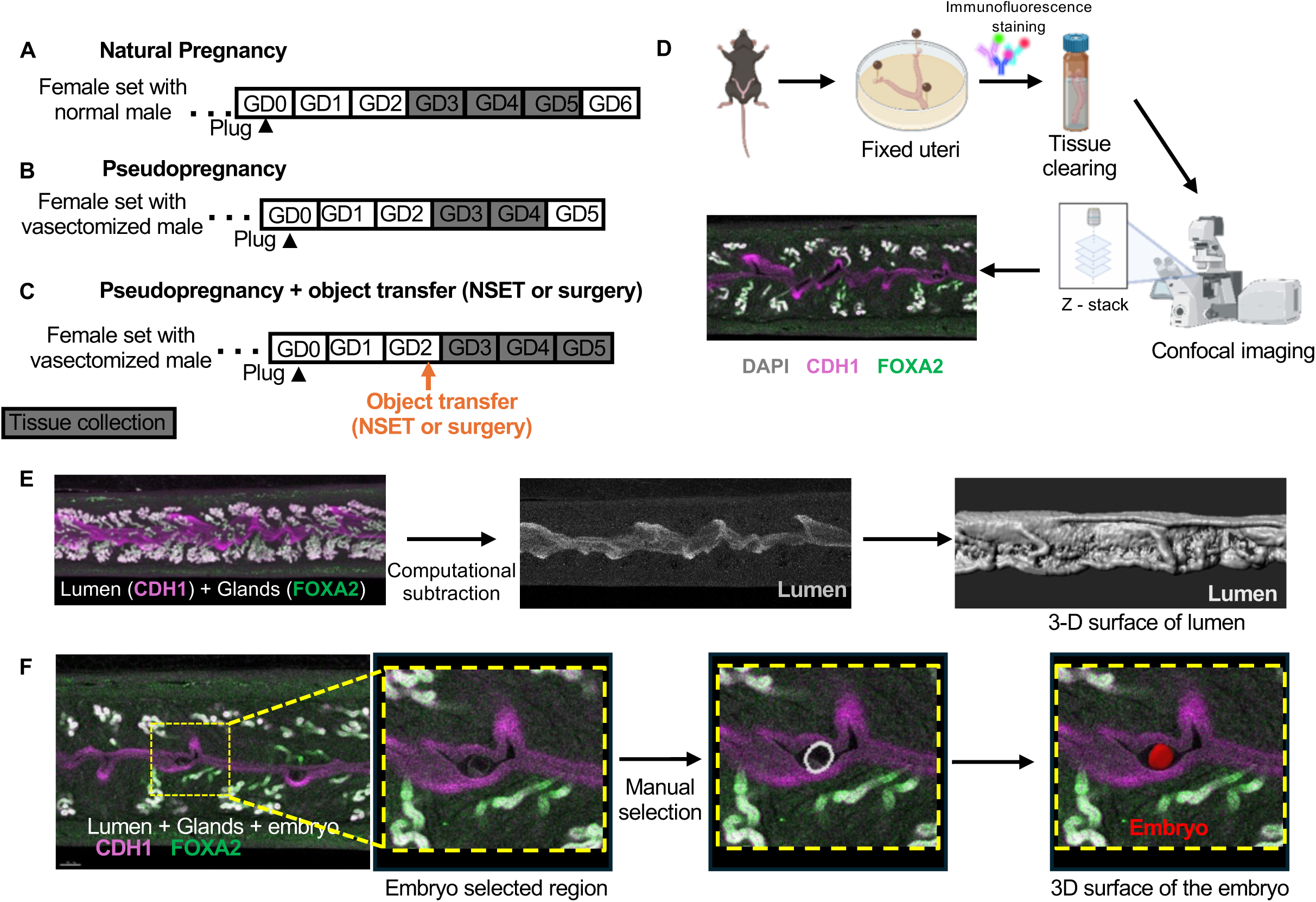
Mouse models and methods used to determine implantation chamber formation is essential for decidualization. **(A-C)** Schematic of mouse model in a natural pregnancy (A), pseudopregnancy (B) and pseudopregnancy + object transfer using NSET or surgery. (D) Illustration of tissue fixation, whole mount immunofluorescence, and imaging protocol. (E-F) Process of 3D reconstruction of the lumen (E) and objects within the lumen (F) using Imaris v9.2.1. Arrowhead is GD0.5 when the plug was identified. Orange arrow indicates the NSET or surgery time. NSET – Non-surgical Embryo Transfer method.

**Supplementary Figure 2.**
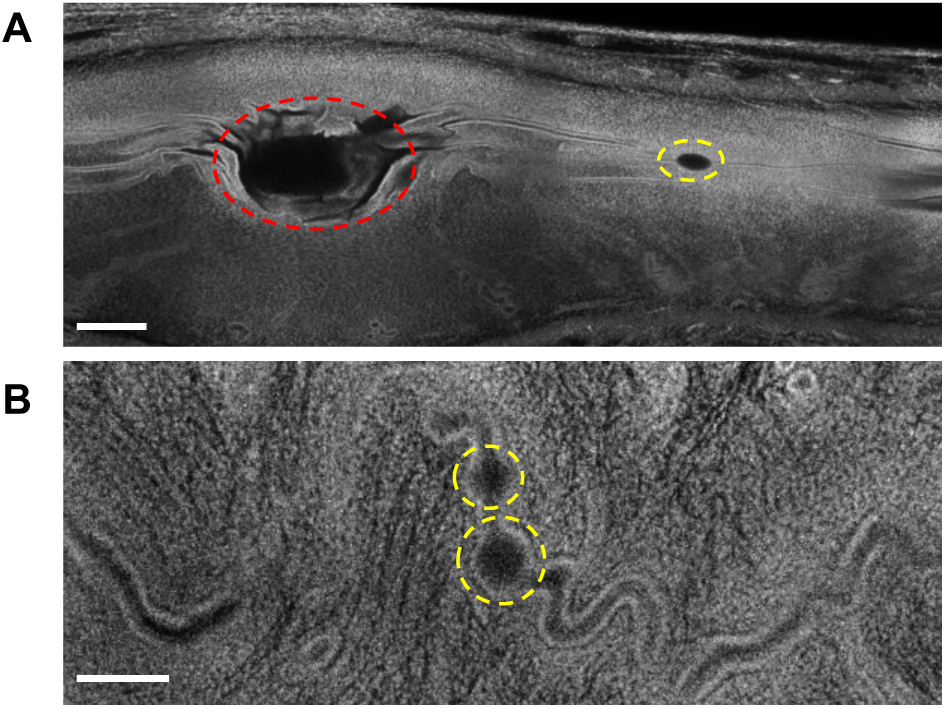
Oil droplets and ConA-beads do not always form implantation chambers. **(A-B)** Representative confocal images showing variable chamber formation at GD4.5: a large oil droplet (red circle) initiates chamber formation, whereas a small oil droplet (yellow circle) does not form a chamber **(A),** two ConA-beads adjacent to each other (yellow circle) that failed to form a chamber **(B)**. Scale bars: **A:** 400 µm, **B:** 100 µm.

**Supplementary Figure 3.**
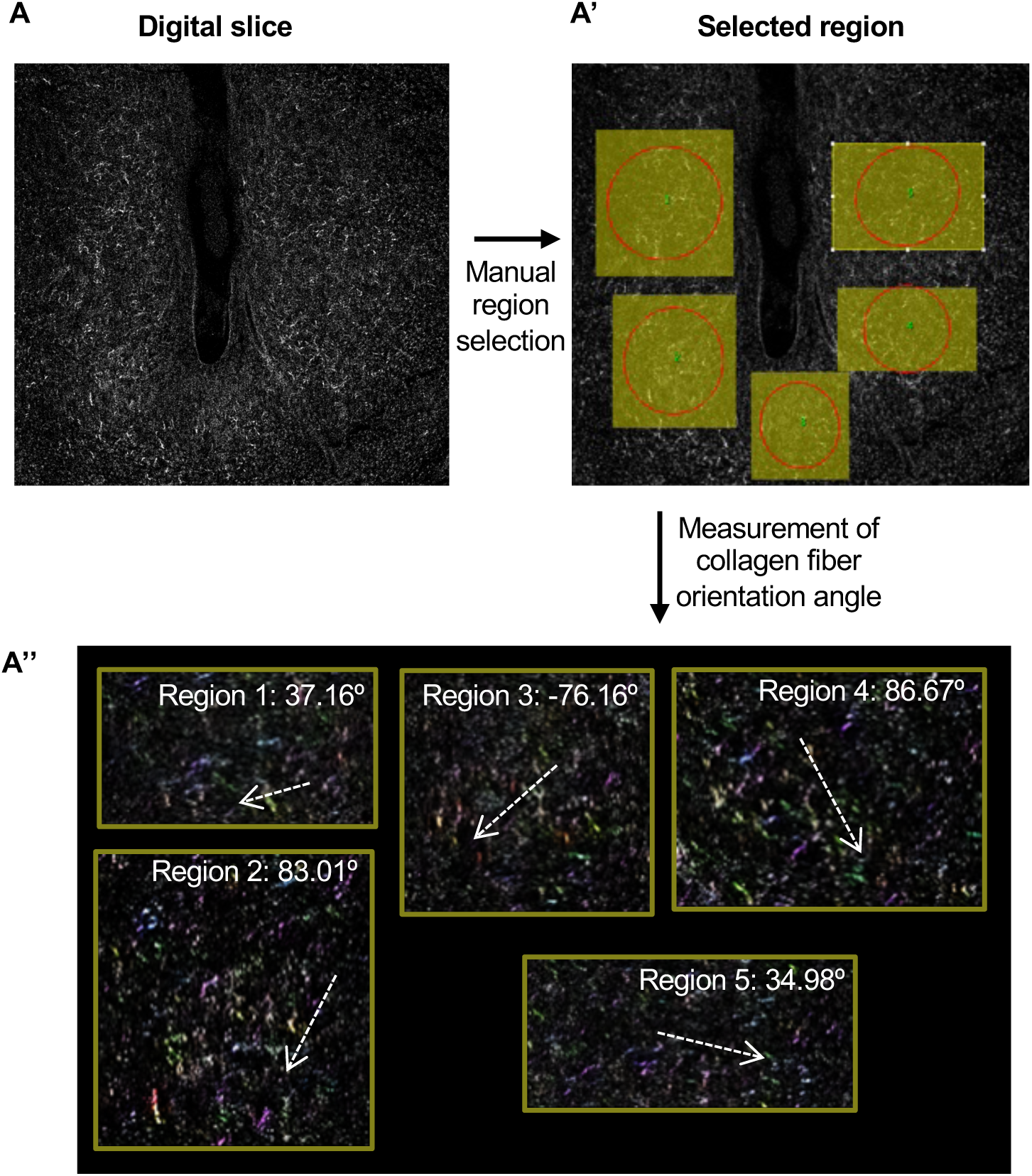
Method for determining fiber orientation from SHG images using the OrientationJ plugin in ImageJ. **(A-A’’)** Pipeline for quantifying collagen fiber angles using OrientationJ involves the following steps: uploading SHG images and converting them to 8-bit format **(A)**, manually selecting five regions surrounding the chamber **(A’)**, and measuring the average angle of collagen fibers in the selected regions **(A’’)**. White arrows indicate the direction of collagen fibers in each region, SHG – Second Harmonic Generation.

## Notes

### Competing Interest Statement

The authors have declared no competing interest.

